# Real-time optimization to enhance noninvasive cortical excitability assessment in the human dorsolateral prefrontal cortex

**DOI:** 10.1101/2024.05.29.596317

**Authors:** Sara Parmigiani, Christopher C. Cline, Manjima Sarkar, Lily Forman, Jade Truong, Jessica M. Ross, Juha Gogulski, Corey J. Keller

## Abstract

**Objective:** We currently lack a robust noninvasive method to measure prefrontal excitability in humans. Concurrent TMS and EEG in the prefrontal cortex is usually confounded by artifacts. Here we asked if real-time optimization could reduce artifacts and enhance a TMS-EEG measure of left prefrontal excitability.

**Methods:** This closed-loop optimization procedure adjusts left dlPFC TMS coil location, angle, and intensity in real-time based on the EEG response to TMS. Our outcome measure was the left prefrontal early (20-60 ms) and local TMS-evoked potential (EL-TEP).

**Results:** In 18 healthy participants, this optimization of coil angle and brain target significantly reduced artifacts by 63% and, when combined with an increase in intensity, increased EL-TEP magnitude by 75% compared to a non-optimized approach.

**Conclusions:** Real-time optimization of TMS parameters during dlPFC stimulation can enhance the EL-TEP.

**Significance:** Enhancing our ability to measure prefrontal excitability is important for monitoring pathological states and treatment response.

**Highlights:**

- We developed a real-time closed-loop optimization procedure to obtain high amplitude early local TEPs (EL-TEPs) from dlPFC TMS.
- Sequential optimization of coil angle and brain target reduced artifacts by 63%.
- Sequential optimization of coil angle, brain target, and intensity increased EL-TEP amplitude by 75%.

## 1. Introduction

Brain stimulation treatments for neuropsychiatric disorders can be highly effective, but the mechanisms by which brain changes elicit clinical response are unclear (Ferrarelli and Phillips, 2021; Lefaucheur et al., 2020). Achieving this understanding will lead to more targeted, personalized treatments, likely translating to improved outcomes. Heavily implicated in neuropsychiatric disorders is the dorsolateral prefrontal cortex (dlPFC), lying at the nexus of multiple brain networks and accessible with noninvasive brain stimulation. Studying dlPFC excitability noninvasively can enhance our understanding of the pathophysiology of these disorders and the brain changes induced by our treatments. To date, however, dlPFC excitability and treatment-related changes have been difficult to probe noninvasively, greatly limiting scientific and clinical progress.

One way to address this challenge is through transcranial magnetic stimulation (TMS) coupled with electroencephalography (TMS-EEG), which provides a causal readout of brain excitability (Ferrarelli and Phillips, 2021; Lioumis et al., 2009). TMS-EEG has provided important insights into the physiology and pathophysiology of sleep (Massimini et al., 2007, 2005), consciousness (Casali et al., 2013, 2010; Edlow et al., 2023; Massimini et al., 2012; Napolitani et al., 2014; Sarasso et al., 2014; Sinitsyn et al., 2020), stroke (Russo et al., 2022; Sarasso et al., 2020), and psychiatric disorders (Donati et al., 2023, 2021; Ferrarelli and Phillips, 2021). However, the majority of these studies measure TMS-EEG in regions where muscle artifact is minimal. In contrast, applying TMS to the dlPFC induces large amplitude muscle artifacts that can obscure the TMS-evoked potential (TEP). This study introduces a new method to reduce muscle artifacts in an effort to enhance the signal-to-noise of prefrontal TEPs.

The TEP is derived from EEG after averaging the response to multiple single TMS pulses (Ahn and Fröhlich, 2021). The early local TEP, or EL-TEP, is a short-latency (20-60 ms) multi-phasic response recorded over the stimulation target (*i.e.*, directly underneath the TMS coil) that occurs prior to large amplitude sensory EEG (Ahn and Fröhlich, 2021; Gogulski et al., 2023b, 2023c; Parmigiani et al., 2023; Ross et al., 2023). Also known as the ‘P20’ and ‘N40’ complex, EL-TEPs may reflect a combination of GABAergic inhibition and NMDA receptor-mediated glutamatergic excitation (Ahn and Fröhlich, 2021; Belardinelli et al., 2021; Hill et al., 2016), and their size has been shown to correlate with depression severity (Voineskos et al., 2019) and predict the clinical response to TMS treatment of depression (Eshel et al., 2020). Intracranial data of similar early potentials induced by single pulse electrical stimulation and TMS support that the EL-TEP is likely to be a measure of cortical excitability (Huang et al., 2019; Keller et al., 2018; Wang et al., 2022). The EL-TEP has been -characterized in multiple brain regions, by our group and others (Ahn and Fröhlich, 2021; Gogulski et al., 2023a; Hill et al., 2016). However, it is difficult to separate the EL-TEP from time-locked evoked muscle artifacts (referred to hereafter as *early artifacts*) that can often last up to 40 ms after applying TMS pulses to the dlPFC, thus confounding interpretation (Mutanen et al., 2013; Rogasch et al., 2013). Offline procedures to clean these artifacts are effective for TMS to brain targets that evoke smaller muscle artifacts (Cline et al., 2021; Korhonen et al., 2011; Mäki and Ilmoniemi, 2011; Mutanen et al., 2018; Rogasch et al., 2017, 2014; Salo et al., 2020; Vafidis et al., 2019; Wu et al., 2018). However, for brain targets that evoke large muscle artifacts, including the dlPFC, minimization during data collection may be highly beneficial, largely due to the difficulty of offline artifact removal. To date, no standardized real-time optimization procedure has been implemented to reduce large-amplitude early muscle-related artifacts like those induced after dlPFC stimulation.

To better understand the relationship between the early artifact and EL-TEPs after dlPFC TMS, we recently applied single pulse TMS to six targets and two angles within the dlPFC (Gogulski et al., 2023b). After offline artifact removal, we observed that TMS to the anterolateral dlPFC at a group level evoked larger early artifacts and smaller EL-TEPs compared to the posteromedial dlPFC. We also observed significant inter-subject variability with respect to an individual’s retrospectively-identified optimal TMS target and coil angle that minimized early artifacts and maximized EL-TEPs. We build on this work here, asking if a real-time adaptive approach for minimizing early artifact can further enhance EL-TEPs.

Current TMS-EEG procedures do not typically involve real-time optimization (Varone et al., 2021). Adjusting coil angle, location, and stimulation intensity in real-time could significantly enhance the TEP. Indeed, a toolbox for real-time TEP monitoring was recently released, with examples on how to optimize experimental approaches (Casarotto et al., 2022). While an important advance, this approach did not delineate a standardized, stepwise, and generalizable procedure to optimize TEPs, and avoided brain regions with large early artifacts. Furthermore, this work focused on real-time TEP visualization, which typically requires averaging >20 pulses to resolve this signal, and often more for regions with large early artifacts (Casarotto et al., 2022; Mutanen et al., 2013). As a result, this method is time consuming, taking 1-2 minutes per parameter combination (>20 trials at ∼3 second inter-pulse interval). With a large parameter space (coil angle, location, intensity), this method may not be practical to use in every TMS-EEG experiment. In contrast, focusing optimization on early artifact may be beneficial because only a single TMS pulse is typically sufficient to quantify early artifacts due to their amplitude, time-course, and topography.

We thus developed and evaluated a real-time optimization procedure to minimize early artifacts and increase the signal-to-noise of EL-TEPs. We hypothesized that by reducing early artifact through coil angle, location, and intensity optimization, we could increase EL-TEP amplitude after dlPFC TMS. We found that in the posterior-medial dlPFC, optimizing angle and location in real-time could reduce early artifact by 63% at low intensity, and when combined with an increase in intensity, increase EL-TEP amplitude by 75% compared to standard procedures. Angle, location, and intensity optimization in combination provided the best enhancement of the EL-TEP. Future work will also include development of fully automated optimization approaches and testing generalizability to other brain regions and populations, leading to tools to efficiently monitor cortical excitability in healthy and pathological brain states and during treatments.

## 2. Methods

### 2.1. Participants

18 healthy participants (22-60 years old, mean=40.61, SD=12.20, six female) provided written informed consent under a protocol approved by the Stanford University Institutional Review Board. All participants completed an online screening questionnaire with inclusion criteria as follows: aged 18-65, fluent in English, able to travel to the study site, and fully vaccinated against COVID-19. Participants were also required to complete the Quick Inventory of Depressive Symptomatology (16-item, QIDS) self-report questionnaire and were excluded if they scored 11 or higher, indicating moderate or more severe depression (Rush et al., 2003). Additional criteria for exclusion included a lifetime history of psychiatric or neurological disorder, substance abuse or dependence in the past month, recent heart attack (<3 months), pregnancy, the presence of absolute contraindications for rTMS (Rossi et al., 2011, 2009), or history of psychotropic medication use. All eligible participants were scheduled for two study visits on separate days to first obtain structural MRI of their brain and then complete a TMS-EEG session. All enrolled participants completed the MRI pre-examination screening form provided by Richard M. Lucas Center for Imaging at Stanford University. All 18 participants completed at least one of the optimization steps and thus were included in the linear mixed effects model analyses, 17 (22-60 years old, mean=41.1, SD=12.7 years old, six female) of which completed all the TMS conditions, and thus were included in all statistical analyses. A table with additional demographic information for participants is available in Supplementary Materials (Table S1).

### 2.2. Transcranial magnetic stimulation

Single-pulse TMS was delivered using a hand-held MagVenture C-B60 figure-of-eight coil with a MagPro X100 stimulator (MagVenture, Denmark). To estimate resting motor threshold (rMT), biphasic single pulses of TMS were delivered to the hand region of the left M1 with the coil held tangentially to the scalp and at 45° from midline. rMT was determined to be the minimum intensity that elicited a visible twitch in relaxed FDI in ≥ 5 out of 10 pulses (Rossini et al., 1994). The optimal motor hotspot was defined as the coil position from which TMS produced the largest and most consistent visible twitch in the relaxed right first dorsal interosseous muscle (FDI).

### 2.3. Target selection

To better understand the effect of targeted dlPFC subregion on EL-TEP optimization, we *a priori* divided the dlPFC into posterior and anterior portions, each containing clinically relevant TMS treatment targets for depression (Gogulski et al., 2023b; Weigand et al., 2018). In the posterior dlPFC, our initial *non-optimized* target was based on a group-level defined treatment location (Weigand et al., 2018) originally 5.5 cm anterior from M1 defined in MNI space (x = -32, y = 26, z = 52). As in our previous work (Gogulski et al. 2023b), anatomical head models were created using SimNIBS headreco (Nielsen et al., 2018), and each MNI target was transformed into an individual’s native MRI space by a nonlinear inverse deformation field (generated using SPM12), with a planned coil orientation generated to position the coil perpendicular to the nearest point on the scalp mesh.

### 2.4. Neuronavigation

Tracking of coil position relative to the subject’s head and MRI data was performed with our custom neuronavigation software, NaviNIBS (manuscript in preparation). Six degree-of-freedom (DOF) poses of the stimulation coil, participant’s head, and stylus tool were tracked via passive infrared optical markers and a Polaris Vega ST camera system (NDI, Waterloo, Ontario, Canada). Registration of participant head tracker position was performed based on fiducials at the nasion and near the left and right preauricular points, with scalp-based refinement. Coil position was visualized relative to the gray-matter and scalp surfaces generated by SimNIBS, with real-time metrics of coil alignment distance and angle offsets to facilitate the optimization procedures described below.

### 2.5. Electroencephalography

EEG data were recorded with a 64-channel TMS-compatible EEG cap with active electrodes (ActiCap, Brain Products GmbH, Germany) and a TMS-compatible amplifier (ActiCHamp plus, Brain products GmbH, Germany) using an acquisition rate of 25 kHz (Mancuso et al., 2021). Electrodes were arranged in a standard montage labeled according to the extended 10-20 international system (Mecarelli, 2019). Electrodes were referenced online to Cz and impedances were generally kept below 5 kΩ, with frequent impedance checks. White noise was delivered through earbuds for the purpose of auditory masking and additional noise damping was provided by passive earmuffs, as previously described (Ross et al., 2022).

### 2.6. Real-time signal processing pipeline

We used a custom Python-based software to allow real-time visualization of TMS-evoked EEG responses. Similar functionality is available with other real-time EEG monitoring software (Casarotto et al., 2022). Our pipeline included the following components:

1. Real-time raw data streaming from our EEG amplifier via the Brain Products RDA protocol (github.com/brain-products/LSL-BrainVisionRDA) and labstreaminglayer (github.com/sccn/labstreaminglayer).
2. Epoch extraction from -500 to 500 ms around each TMS pulse.
3. Interpolation of primary artifact timespan from -5 to 10 ms around each pulse.
4. Downsampling to 1 kHz.
5. Baseline subtraction, using a baseline period of -500 to -50 ms before each pulse.
6. Bad channel rejection and interpolation of any manually-identified bad channels.
7. High-pass filtering above 5 Hz.
8. Notch filtering at 60 Hz.
9. Low-pass filtering below 50 Hz.
10. Average re-referencing.
11. Extraction of peak-to-peak global mean field amplitude (GMFA) in a 10-20 ms time window after each pulse, i.e. a short-latency window beginning immediately after interpolate of primary TMS-evoked artifact.
12. Visualization of recent history of per-trial extracted GMFA features updated after each pulse.
13. Visualization of evoked response waveforms and topographies, updated after each pulse.

### 2.7. Closed-loop optimization procedure

Through a stepwise real-time procedure, we aimed to enhance the quality of prefrontal EL-TEPs by minimizing early-latency TMS-evoked artifacts. Using the pipeline described in the previous section to quantify TMS-evoked artifact in a 10-20 ms period after each pulse, we applied a real-time optimization algorithm consisting of the following steps : (1) *coil angle optimization*: keeping the coil centered at an initial pre-planned target and tangential to the scalp, sampling a range of angles; (2) *location optimization*: using a fixed angle identified in step 1 and sampling multiple locations in a region near the initial target; (3) *coil angle refinement*: varying angle in a smaller range centered on the angle identified in step 1 at the location target identified in step 2; and (4) *stimulation intensity increase:* increasing intensity from the initial 110% rMT up to 140% rMT, when tolerable, applied at the optimal location and angle identified in step 3. See below for more details of each of these steps and related data acquisition.

**(1) *Coil angle optimization*** (Fig 1A). Single TMS pulses were delivered at our predefined MNI location, keeping the coil tangential to the scalp while modifying angle in the coil’s horizontal plane (Fig 1A and Fig 2A-C). During this angle optimization procedure, we identified the coil angle that produced the smallest early artifact (defined as amplitude of the 12-20 ms GMFA). Angle differences between adjacent samples were less than ten degrees, with a higher density of samples near local minima in early artifact, and with angle constraints determined by limitations in neuronavigation camera tracking and TMS coil cable length. Stimulation intensity was 110% rMT (stimulation intensities for all participants are shown in Table S3). See Figure 2C (group) and Fig S1 (each participant) for the relationship between coil angle and size of the early artifact. The angle that produced the smallest early artifact was chosen for the subsequent location optimization step (Fig 2A, C). Averaged across participants, this step took 9.8 ± 3.8 (mean ± SD) minutes.
**(2) *Location optimization*** (Fig 1B). While maintaining the optimal coil angle determined in the previous step and orienting the coil tangential to the scalp, the coil was moved within a circular region with a radius of 1.5 cm around the original, non-optimized target (Fig 1B). 1.5 cm was chosen to balance several considerations, including a large enough region to observe variation in artifact and close enough to the initial target to be functionally and clinically relevant for depression treatment (Fox et al., 2012; Gogulski et al., 2023b; Herwig et al., 2003; Weigand et al., 2018). As in step 1, in this location optimization step, the location that produced the smallest artifact was identified. This step took on average 6.7 (± 2.7) minutes.
**(3) *Angle refinement*** (Fig 1C). To check whether the optimized angle from step 1 was still the best angle at the optimized location from step 2, we introduced an additional angle refinement step, in which we applied single TMS pulses at the optimal location but with small variations in coil angle around the step 1 determined angle. This step took 5.7 (± 3.3) minutes.
**(4) *Intensity increase*** (Fig 1C). After optimizing coil orientation at 110% rMT in the previous steps, we increased stimulation intensity up to 140% rMT or an intermediate intensity as limited by participant tolerability (see Table S3 for participants’ highest tolerated intensities).

**Fig 1.**
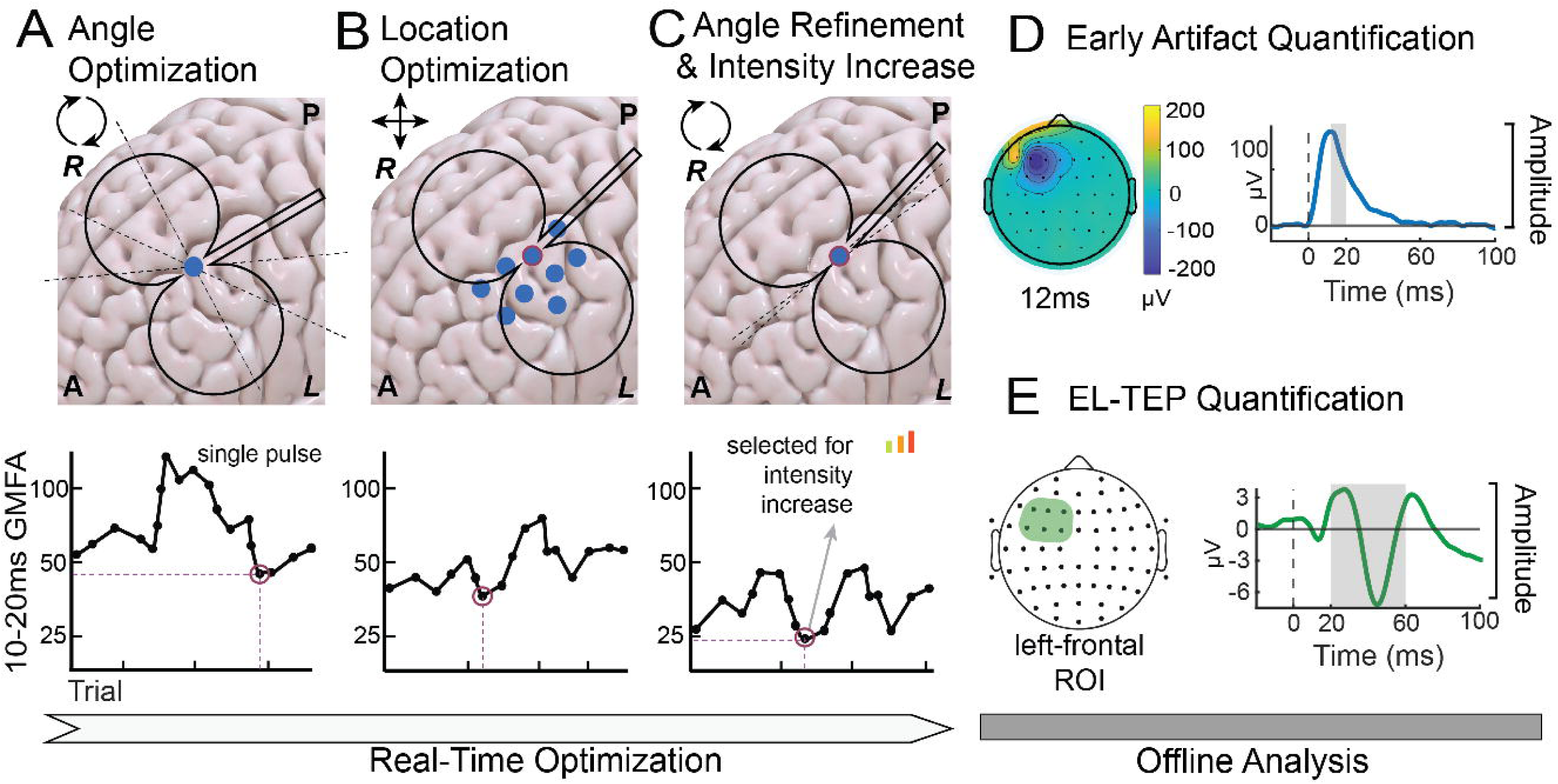
Workflow. **A-C**) *Real-time procedure for reducing early artifact.* A) Angle optimization, beginning at 45 degrees with respect to the midline at the MNI coordinate defined, non-optimized, target (x = -32, y = 26, z = 52). B) Spatial search within 1.5 cm radius around initial target at optimal angle resulting from A. C) Angle adjustment at optimal stimulation site. Once the angle that induced minimal early artifact was determined, stimulation was delivered at multiple intensities between 110-140% rMT to study the effect of intensity on the TEP. On the top row, schematic brains are depicted with a coil over the targets. Coordinates (Anterior, Posterior, Left, and Right) are indicated with their initials. On the bottom row, per-trial GMFA values are visible in the charts, with each dot representing an example trial. For each step, the lowest value is circled and used as coordinates for the following step. **D-E**) *Offline analysis.* D) Selected time window for quantifying the early artifact (GMFA 12-20 ms after TMS pulse, outlined in gray) and an example of topography of early artifact at 12 ms after minimal preprocessing (see 3.1 and 3.2 for definition of minimal preprocessing). E) Region of interest (left prefrontal) and time course (20-60 ms, in gray) of the EL-TEP.

**Fig 2.**
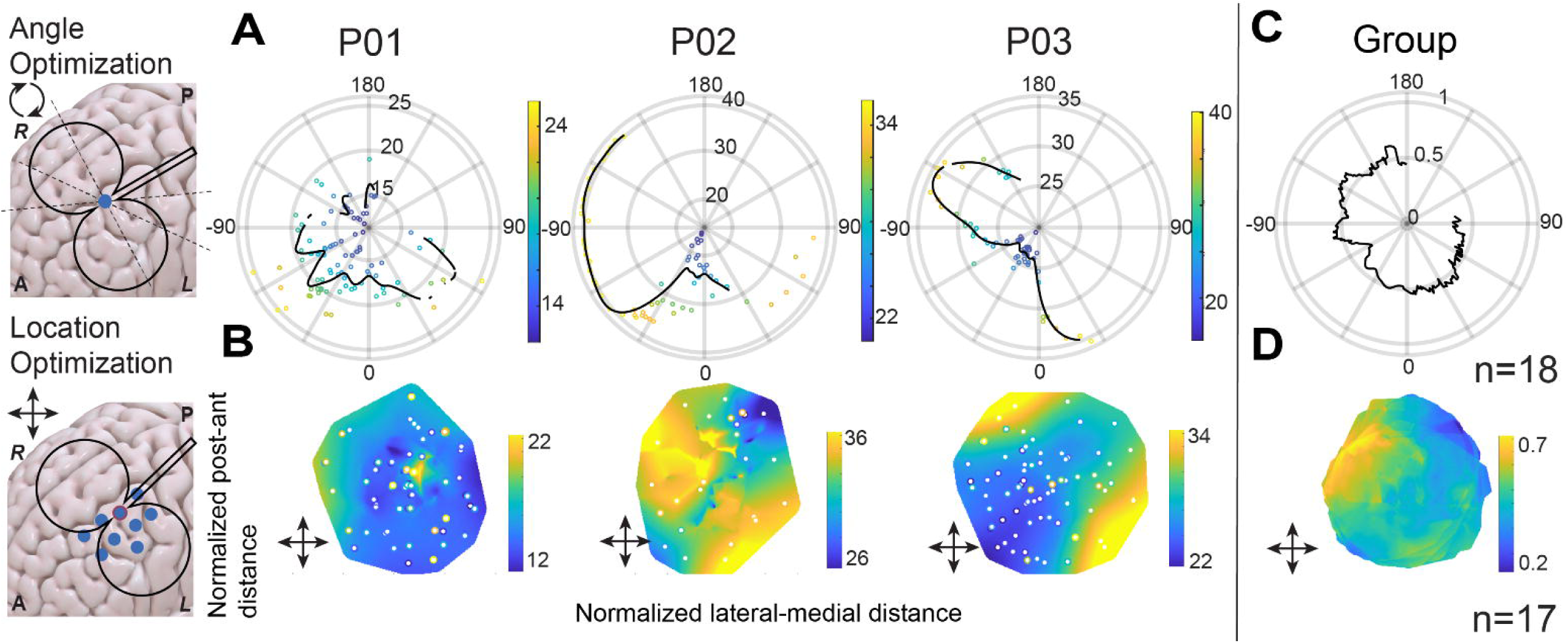
Real-time optimization reduces the early artifact. Three example participants (A, B) and group data averages (C, D). A) Coil angle search procedure for three different participants (P01, P02, P03). GMFA of the early artifact (in dBμV at 12-20 ms after TMS) are shown in relation with angles from the midline - the lower the artifact, the closer to the center of the radial representation. B) Spatial search for location optimization in the same three participants. GMFA of early artifacts are shown in relation with normalized lateral-medial distance (x-axes) and normalized posterior-anterior distance (y-axes). C) Group level effect of angle optimization (N=18). D) Group-level effect of spatial search for location optimization (N=17).

### 2.8. Data collection

To compare offline the TEPs across optimization conditions, we collected blocks of 150 single TMS pulses in six conditions enumerated below. Since coil angle, location, and intensity were kept constant within each of these blocks, we refer to these blocks as “static” conditions. To examine the effect of optimizing coil location and angle, we considered three conditions: i) a *non-optimized* location and orientation at the initial target with 45 degree coil placement with respect to the midline (as is common practice, without any optimization), ii) the initial target at the *optimal angle* determined from the step 1 angle optimization, and iii) the *optimal angle and location,* determined from steps 1-3.

To investigate the effect of stimulation intensity and interaction with location and orientation optimization, each of these conditions was tested at two stimulation intensities: at 110% rMT and at a higher intensity of 140% rMT when tolerable for the participants and lower if not tolerable (Table S3 for participants’ highest tolerated intensities). The rationale of this range choice relies on the possibility to investigate suprathreshold intensities that are also commonly used in clinical settings. Our static conditions, each with 150 pulses in each stimulation configuration, can be therefore described as follows: i) *non-optimized* (45-degree angle and initial location)*, 110% rMT*; ii) *angle optimized, 110% rMT*; iii) *angle and location optimized, 110% rMT*; iv) *non-optimized*, *high intensity*; v) *angle optimized*, *high intensity,* and vi) *angle and location optimized*, *high intensity*.

## 3. Data Analyses

### 3.1. Preprocessing of TMS-EEG data

For each recorded block of 150 pulses, data were first preprocessed as follows, using version 2.0 of the fully automated AARATEP pipeline. Epochs were extracted from 800 ms before to 1000 ms after each TMS pulse. Data between 2 ms before to 12 ms after each pulse were replaced with values interpolated by autoregressive extrapolation and blending (Cline et al., 2021), downsampled to 1 kHz, and baseline-corrected based on mean values between 500 to 10 ms before the pulse. Epochs were then high-pass filtered above 1 Hz with a modified filtering approach (see Cline et al 2021 for details) to reduce the spread of pulse-related artifact into baseline time periods. Bad channels were rejected via quantified noise thresholds and replaced with spatially interpolated values. Eye blink artifacts were attenuated by a dedicated round of independent component analysis (ICA) and eye-specific component labeling and rejection using ICLabel (Pion-Tonachini et al., 2019), a modification from the original AARATEP pipeline introduced in version 2.0. Various non-neuronal noise sources were attenuated with SOUND (Mutanen et al., 2018). Early artifacts were reduced via a specialized decay fitting and removal procedure (Cline et al., 2021). Line noise was attenuated with a bandstop filter between 58-62 Hz. Additional artifacts were attenuated with a second stage of ICA and ICLabel labeling and rejection, with rejection criteria targeted at removing any clearly non-neural signals. Data were again interpolated between -2 and 12 ms with autoregressive extrapolation and blending, low-pass filtered below 100 Hz, and average re-referenced. For complete details of the pipeline implementation, see Cline et al. 2021, and source code at <u>github.com/chriscline/AARATEPPipeline</u>.

For analysis of early artifacts recorded during optimization and in 150-pulse static blocks, a separate “minimal” processing pipeline was used, with a subset of steps from the full AARATEP preprocessing pipeline described above. Specifically, epochs were extracted from 800 ms before to 1000 ms after each TMS pulse, down sampled to 1 kHz without filtering (to prevent dispersion of primary pulse artifact), and baseline-corrected. Data between 2 ms before to 12 ms after each pulse were interpolated and bad channels were rejected and interpolated as described above. Epochs were band-pass filtered between 1 Hz and 100 Hz with the same modified filtering approach mentioned above, and average re-referenced.

### 3.2. Quantification of the early artifacts and the EL-TEP

The two main features extracted from the EEG response to single pulses of TMS to the dlPFC were i) *early artifact* defined as the mean GMFA from 12 to 20 ms from the minimally processed data and based on ICA topography and spectral features consisting primarily of evoked craniofacial muscle twitches and (Casarotto et al., 2022; Mutanen et al., 2013) ii) the prefrontal *EL-TEP*, defined as the min-to-max amplitude difference within 20 to 60 ms after the TMS pulse at electrodes over the site of stimulation from the fully processed data (Gogulski et al., 2023a). For this prefrontal EL-TEP, we used a region of interest (ROI) consisting of six electrodes (F5, F3, F1, FC5, FC3, FC1; see Fig 3). These electrodes were chosen to broadly cover the left prefrontal cortex and have been selected previously to quantify the EL-TEP (Gogulski et al., 2023b, 2023a). To explore the effect of real-time optimization across non-local distributed brain regions, we also quantified early (but non-local) TEPs in the 20-60 ms window in a contralateral *right frontal* ROI (F2, F4, F6, FC2, FC4, FC6) and in a *left parietal* ROI (CP3, CP1, P5, P3, P1, PO3; see Fig 4). Per-channel averages were calculated with a trimmed mean rejecting the most extreme 10% of values across trials at each time point.

**Fig 3.**
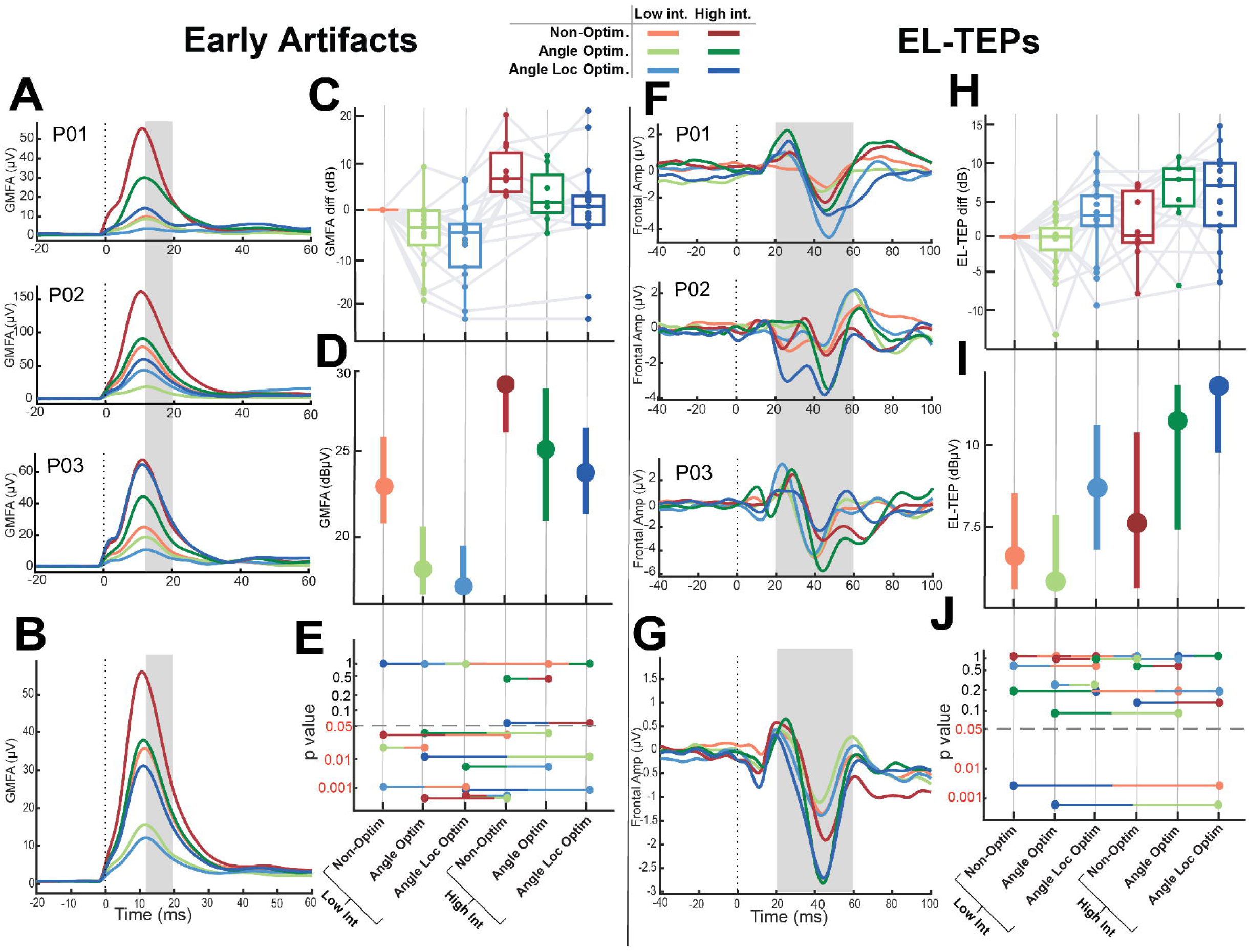
Real-time optimization reduces the early artifact and enhances the early local TMS-evoked potential. A and F) Individual participants. A) Early artifacts in three participants. Plots show the GMFA of minimally preprocessed data (see Methods). Gray bar denotes the time window for GMFA calculation (12-20ms); and F) EL-TEPs in three example participants. Gray bar denotes the time window for EL-TEP calculation (20-60ms). B and G) Group-level results (N=18), B) GMFA traces across conditions for early artifact; and G) signal across conditions for EL-TEPS. Artifact and EL-TEPs across conditions boxplots with individual participants connecting dots are in C) for GMFA early artifact and J) for EL-TEPs. C-I) Quantification of group results based on LME results and EMMeans predictions comparisons. Grids are reported in D) for GMFA early artifact and H) EL-TEPs. All p-values from the EMMeans comparisons are in E) GMFA early artifact and I) EL-TEPS.

**Fig 4.**
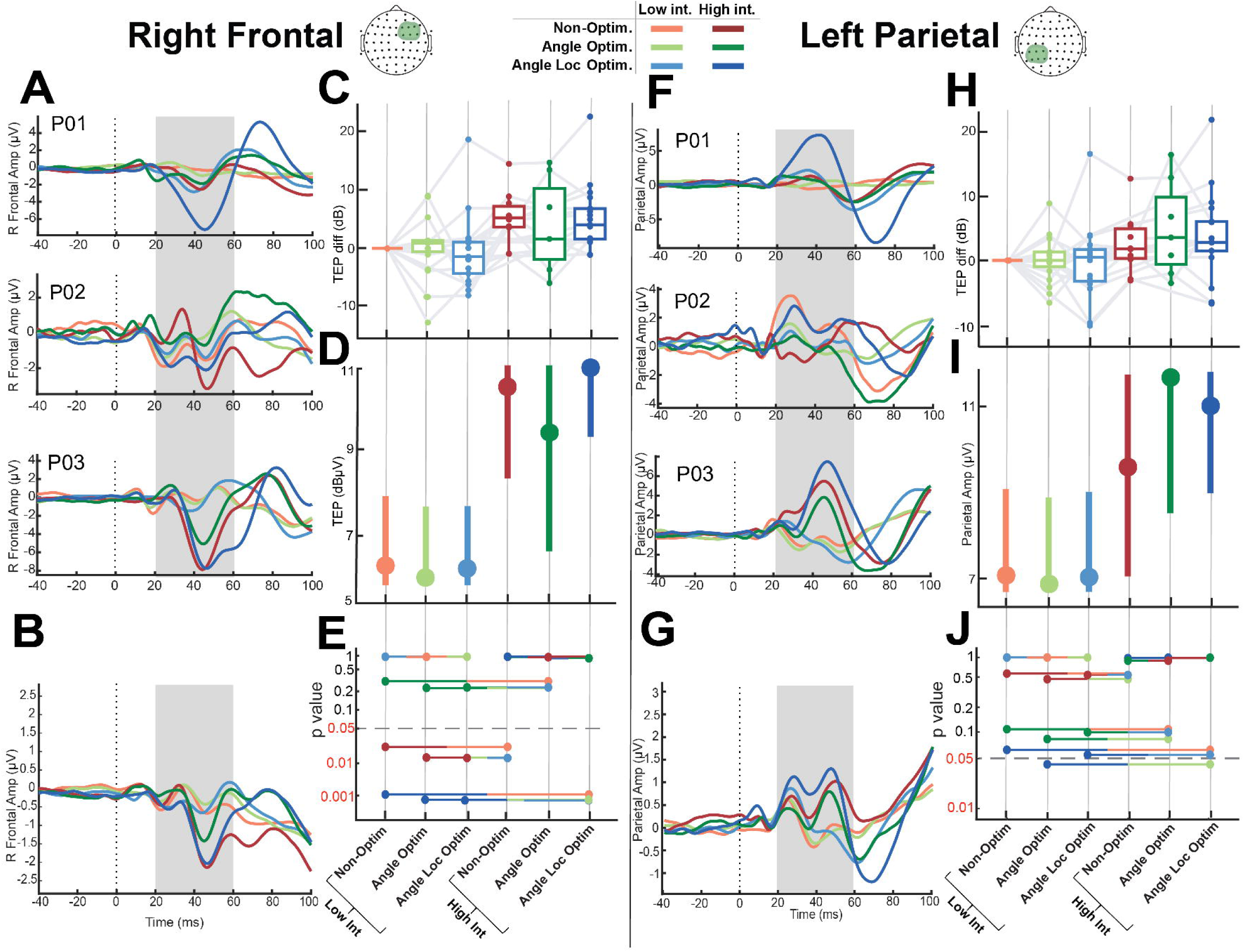
Real-time optimization reduces the early artifact and enhances the early local TMS-evoked potential of distal cortical areas in a dose-dependent manner. A and F) Single participant examples of EL-TEPs in A) contralateral right frontal ROI and F) left parietal ROI. The early TEP time window (20-60 ms) is highlighted in gray. B and G) Group-level results (N=18), early TEPs across conditions boxplots with individual participants connecting dots are in C) for contralateral right frontal ROI and J) for left parietal ROI. C-I) Quantification of group results based on LME results and EMMeans predictions comparisons. Grids are reported in D) for contralateral right frontal ROI and H) for left parietal ROI. All p-values from the EMMeans comparisons are in E) contralateral right frontal ROI and I) left parietal ROI.

### 3.3. Statistical analyses

All statistical analyses were performed in R version 4.2.3 (R Core Team, 2013). Analyses were performed to test if the real-time optimization process i) reduced early artifacts and ii) enhanced EL-TEP amplitudes. Before evaluating the statistical measures, each subset of data was tested for normality (Shapiro and Wilk, 1965). We first performed paired samples t-tests on participants who completed the optimization procedure and had balanced data across conditions (*n*=17, excluding P08), focusing on contrasts related to our main hypotheses. The conditions compared were i) early artifacts in the *non-optimized* condition at 110% rMT vs. fully optimized stimulation (*optimized angle* and *location* at 110% of rMT); and ii) the *non-optimized* EL-TEPs at 110% rMT vs. fully optimized EL-TEPs (*optimized angle* and *location* at the *highest intensity*). Then, to account for the unbalanced sample sizes across conditions, we used linear mixed-effects models (Bates et al., 2015; Fox, 2008; Luke, 2017) (LME; n=18) and compared the effect of different optimization steps on early artifacts and EL-TEPs, followed by *post hoc* comparisons using Tukey’s adjustment for pairwise comparisons. In our model, we specified the *optimization steps* of the procedure as fixed effects to assess their direct impact, while including *participants* as a random intercept term to account for individual differences. This LME approach allowed for modeling of within-subject effects with incomplete data (*i.e.*, that some conditions were only collected in a subset of participants). We computed estimated marginal means (EMMs) (Lenth, 2019) to summarize model values of each fitted LME model at specific factor levels. If data included in an LME are unbalanced, as in our case, the EMMs represent an estimate of the marginal means that we would have observed if the data had been balanced across conditions (Lenth, 2019). Finally, to investigate whether optimization affected the early non-local TEP, we repeated the paired samples t-testing and LME in the right frontal and left parietal ROIs.

## 4. Results

### 4.1. Real-time optimization reduces the TMS-evoked early artifact

We first asked if our optimization algorithm reduced the early TMS-induced artifact as intended. At the individual level, qualitatively some participants demonstrated a robust sensitivity of coil angle to early artifact (*e.g.,* P02, P03 in Fig 2A) while others (P01 in Fig 2A) did not (see Fig S1 for angle sensitivity maps for each participant). Similarly, during location optimization, some participants demonstrated a clear spatially-varying artifact magnitude map (*e.g.*, P03 in Fig 2B) while others (P01, P02 in Fig 2B) did not. At the group level, during optimization we observed a sensitivity of early artifacts to specific angles (Fig 2C) as well as locations (Fig 2D). The *non-optimized 110% rMT* condition resulted in an average early artifact of 26.1 μV (range 1.66-97.33 μV, SD 26.9), while the *angle and location optimized 110% rMT* condition resulted in an average early artifact of 9.6 μV (range 1.8-42.32 μV, SD 10.9), an average reduction of 63.4% (expressed in linear value, μV) in early artifact after optimization. A paired *t*-test of the GMFA expressed in dBμV confirmed that early artifact was significantly higher in the *non-optimized* condition at 110% rMT compared to the *angle and location optimized* condition at 110% rMT (N=17, t(16) = 3.26, p = .0048). Next, we contrasted the estimated marginal means (EMMs) from our LME (N=18) predictions (see Fig 3C for LME, 3D for EMMs comparisons, and 3E and Table S5 for p-values), adjusting for multiple comparisons, and found significant differences between several conditions. Of particular interest, i) *angle optimized* early artifact at 110% rMT was significantly smaller than *non-optimized* early artifact at 110% rMT (5.42 dB difference, t(65.0) = 3.31, p = .0181); and ii) *angle and location optimized* artifact at 110% rMT was significantly smaller than *non-optimized* artifact at 110% rMT (6.67 dB difference, t(65.1) = 4.12, p = .0012).

### 4.2. Real-time optimization enhances the EL-TEP in the dlPFC

Next, we asked if our real-time optimization procedure to minimize early artifact also increased EL-TEP amplitude. At the individual participant level, we qualitatively observed different magnitudes of EL-TEP change with coil *angle optimization*, *angle and location optimization*, and *intensity increase* (Fig 3J, all individual participants in Fig S3). Quantitatively, peak-to-peak amplitude of *non-optimized* EL-TEPs at *110% rMT* were on average 2.5 μV (range 0.8-6.6 μV, SD 1.87), while *angle and location optimized* at *higher intensity* were on average 4.4 μV (range 1.3-9.1 μV, SD 2.34), with an increase of EL-TEPs of 75.5%. A paired t-test indicated that *angle and location optimized* EL-TEPs at the highest intensity were significantly larger than *non-optimized* EL-TEPs at 110% rMT (t(16) = 3.74, p = .00179). LME results (N=18) also showed significant group-level effects of optimization strategies (*coil angle and location*) and intensity on the EL-TEP (Fig 3J). Specifically, we observed that based on contrasts of EMMs (Fig 3H): i) *angle and location optimized* at the highest intensity produced a significantly larger EL-TEP than *non-optimized* at 110% rMT (-5.47 dB difference, t(65.2) = -3.85, p = .0036); and ii) *angle and location optimized* at the highest intensity produced a significantly larger EL-TEP than *angle optimized 110% rMT* only (-6.31 dB difference, t(65.3) = -4.37, p = .0006). No other comparisons demonstrated a significant difference (all p-values shown in Fig 3J and Table S5). However, it is worth noting that EL-TEPs from the *non-optimized highest intensity* were not significantly different from the EL-TEPs from the *non-optimized 110% rMT* condition (-1.14 dB difference, t(66.5) = -0.67, p = .98), suggesting that intensity alone did not have a significant effect on the EL-TEP. Taken together, our results indicate that real-time angle and location optimization combined with an increase in stimulation intensity increased EL-TEP amplitude compared to other strategies.

### 4.3. Real-time optimization of dlPFC TMS affects the distal early TEP in a dose-dependent manner

Next, we asked if and how after left dlPFC TMS real-time optimization affected the early (20-60ms) TEP in the contralateral frontal and ipsilateral parietal ROI. In the right frontal ROI, we observed that right frontal early TEPs after *angle and location optimization* at the *highest intensity* were stronger than right frontal early TEPs *non-optimized* at *110% rMT* (t(16) = 3.91, p = .0012, paired t-test). However, using EMMs from our LME model incorporating all participants, we found significant effects of *intensity* only (N=18, Fig 4D): i) *non-optimized* right frontal early TEPs at highest intensity were significantly larger than *non-optimized* early TEPs at 110% rMT (-4.97 dB difference, t(66.9) = -3.25, p = .0214); ii) *angle and location optimized* right frontal early TEPs at highest intensity were significantly larger than *non-optimized* early TEPs at 110% rMT (-5.34 dB difference, t(65.2) = -4.18, p = .0012); iii) *non-optimized* right frontal early TEPs at the highest intensity were larger than *angle optimized* early TEPs at 110% rMT (-5.31 dB difference, t(67.3) = -3.42, p = .0132); iv) *angle and location optimized* right frontal early TEPs at the highest intensity were larger than *angle optimized* early TEPs at 110% rMT (-5.68 dB difference, t(65.4) = -4.38, p = .0006); v) *non-optimized* early TEPs at the highest intensity were larger than a*ngle and location optimized* right frontal early TEPs at 110% rMT (-5.22 dB difference, t(67.3) = -3.42, p = .0129); and vi) a*ngle and location optimized* right frontal early TEPs at the highest intensity were larger than a*ngle and location optimized* early TEPs at 110% rMT (-5.59 dB difference, t(65.8) = -4.40, p = .0005). P-values are reported graphically in Fig 4E and listed in Table S5. In summary, for a given intensity level, angle or angle and location optimization after left dlPFC TMS produced minimal differences in the right frontal early TEPs. In contrast, higher intensities had a notable impact.

In the left parietal ROI, early TEPs were larger in the *angle and location optimized* condition at highest intensity compared to the *non-optimized* condition at 110% rMT (t(16) = 2.24, p = .0396, paired t-test). However, in our EMM contrasts (N=18) only left parietal early TEPs from a*ngle and location optimized* at the highest intensity was significantly different than those from *angle optimized* at 110% rMT (Fig 4H; - 3.90 dB difference, t(65.5) = -3.01, p = .0410). All p-values are reported graphically in Fig 4J and listed in Table S5.

## 5. Discussion

In the present study, we sought to enhance a measure of cortical excitability by investigating the effects of real-time optimization of TMS angle, location, and intensity on the prefrontal EL-TEP. We hypothesized that reducing early artifacts through real-time adaptive optimization would unmask underlying direct cortical responses to prefrontal TMS, thus increasing the size of observed EL-TEPs. In 18 healthy participants, we observed the following: optimization of TMS coil angle and target reduced early evoked local artifact by 63% (Fig 3B) at an intensity of 110% RMT, and when combined with an increase in intensity increased prefrontal EL-TEPs by 75% (Fig. 3G).

Optimizing coil angle and location alone reduced early artifact compared to the non-optimized condition. While higher stimulation intensities generally invariably resulted in larger early artifacts, the trends in early artifact between no optimization, angle optimization, and angle and location optimization were similar within each intensity level.

Only optimizing coil angle, or angle and location, did not significantly affect EL-TEPs. Only increasing intensity without optimizing coil angle or location also did not significantly affect EL-TEPs. But combining coil angle and location optimization with an increase in intensity significantly increased EL-TEPs, indicating the importance of combining optimization of both coil position and stimulation intensity.

In contrast, non-local TEPs in left parietal and right frontal ROIs were larger at higher intensities, without a strong effect of optimization strategy within each intensity level (Fig 4). This is in line with our hypothesis that local responses were effectively masked by (local) stimulation artifact, and that our artifact minimization procedure enabled observation of these previously masked responses, while not strongly affecting downstream non-local responses elsewhere in the brain.

Together, this work introduces a standardized and relatively fast real-time optimization approach that enhances prefrontal EL-TEPs. This approach should be considered for use in monitoring prefrontal excitability in pathological brain states and during treatments.

### 5.1. Comparison to previous approaches

When stimulating the dlPFC and other regions near craniofacial muscles, large artifacts are common (Mutanen et al., 2013), degrading the ability to measure. high quality prefrontal EL-TEPs. To improve data quality after dlPFC TMS, attention has been given to TMS coil type and shape, dynamic range of the EEG amplifier, wire orientation of the electrodes, and electrode impedances (Casarotto et al., 2022; Mancuso et al., 2024, 2021; Varone et al., 2021). Although careful selection of these parameters can reduce some sources of early artifact, the quality of TEPs in prefrontal areas generally remained low. In early TMS-EEG studies, it was suggested that the coil should be “carefully rotated to yield minimal stimulation artifact” to achieve a clean response (Cracco et al., 1989). While the recent release of software for real-time TEP monitoring (Casarotto et al., 2022) has provided the field a tool for minimizing artifacts, a standardized, stepwise, and generalizable procedure to amplify TEPs has not yet been described. Moreover, existing examples using this toolbox have prioritized real-time TEP visualization which, based on Casarotto and colleagues’ (2022) findings, requires averaging at least 20 pulses to resolve the signal even in brain regions outside the dlPFC with typically weaker early artifacts. Our work introduces for the first time a standardized and adaptive optimization algorithm to tackle the persisting challenges in a region with significant early artifact, enabling improved signal-to-noise measurement for future studies of prefrontal excitability.

### 5.2. Application of EL-TEP optimization

This real-time optimization approach introduced here could have broad applications in understanding physiology and pathophysiology in neurological and psychiatric disorders. The early local response to single pulses of TMS (here and in our former work (Gogulski et al., 2023b, 2023b; Ross et al., 2023) referred to as EL-TEPs) are thought to at least in part index cortical excitability (Belardinelli et al., 2021). Since the dlPFC is a commonly used therapeutic target for TMS treatment and a core region putatively involved in psychiatric disorders such as depression (Ferrarelli and Phillips, 2021; Lefaucheur et al., 2020), our focus has been to evaluate local cortical changes via EL-TEPs in and around the dlPFC.

However, EL-TEPs in the lateral prefrontal cortex are highly likely contaminated by muscle-evoked artifacts, which can significantly confound results and limit interpretability. Our previous work demonstrated post-hoc differences in the amplitude and reliability of the EL-TEP across six dlPFC targets (Gogulski et al., 2023a, 2023b). Here we expanded on this work by using real-time optimization to efficiently explore the effects of varying single-pulse TMS parameters, finding optimal parameters for minimizing early artifacts and thus uncovering EL-TEPs. In the end, higher fidelity measures of prefrontal excitability (EL-TEPs) will allow researchers to confidently sample prefrontal excitability changes after treatments such as repetitive TMS and facilitate the development of more effective, targeted, and personalized treatments.

### 5.3. Limitations and future directions

Despite the promising results reported here, several limitations need to be addressed in future research. This study enrolled healthy participants, and it is unknown how well this optimization algorithm will perform in other populations, such as those with psychiatric disorders. Testing this algorithm in patients is necessary to determine its clinical relevance. The performance of the optimization with different EEG systems and TMS pulse types, coils, and stimulators remains unclear. The current study focused on optimizing TMS parameters to minimize early artifact in real-time and to enhance the EL-TEP offline. Future research should assess the effect of this procedure on other TMS-EEG measures, including later (>100ms) evoked components of the TEP and induced oscillations. The algorithm tested here did not directly adapt stimulation based on the EL-TEP in real-time, but instead minimized early artifact. This type of approach should be contrasted with efforts that manipulate TMS parameters directly based on the EL-TEP amplitude (e.g., Casarotto et al., 2022). For simplicity we employed a sequential optimization approach focusing on angle, location, and intensity without thoroughly investigating their interactions and future work should explore these interactions more exhaustively. This study primarily focused on one dlPFC subregion. Further work is needed to determine if this algorithm can also minimize artifact and enhance EL-TEPs after TMS to other dlPFC subregions including anterior dlPFC, a relevant target for TMS treatment for depression, and other cortical targets. Future work should test this and other approaches on dlPFC and on different brain regions.

Lastly, refining our optimization algorithm by implementing more efficient search procedures and fully automating the protocol could yield faster and potentially more consistent results. The search constraints outlined in the optimization algorithm presented in this work can also be tailored in future studies based on specific requirements. This may involve limiting the duration of optimization or constraining stimulation location to not deviate too far from a clinically-defined target. We used a manual approach to optimize TMS parameters, which took on average 21 minutes (Table S4). Future research will automate this work to standardize further and reduce overall time needed.

## 6. Conclusions

We introduce a real-time TMS-EEG optimization approach that successfully minimized early artifacts and increased the size of prefrontal EL-TEPs. This approach has important implications for investigating and monitoring prefrontal excitability, with applications in studying healthy and diseased brain states and quantifying clinical treatment effects. Future work should include comparing different optimization procedures, testing this algorithm in other brain regions and participant populations, and automation.

## Acknowledgements

We extend our gratitude to all our research participants. We would also like to acknowledge the generous contributions of the members of the Stanford Laboratory for Personalized Neurotherapeutics for helpful feedback on the manuscript and throughout the course of the study. This research was supported by the National Institute of Mental Health under award numbers R01MH126639, R01MH129018, and a Burroughs Wellcome Fund Career Award for Medical Scientists (CJK). JG was supported by personal grants from Orion Research Foundation, the Finnish Medical Foundation and Emil Aaltonen Foundation. JMR was supported by the Department of Veterans Affairs Office of Academic Affiliations Advanced Fellowship Program in Mental Illness Research and Treatment, the Medical Research Service of the Veterans Affairs Palo Alto Health Care System, and the Department of Veterans Affairs Sierra-Pacific Data Science Fellowship.

