## Supplemental Information for "Real-time optimization to enhance noninvasive cortical excitability assessment in the human dorsolateral prefrontal cortex"

*Table S1 - Participant demographics. N = 18. Participants were on average 40.61 years old (SD = 12.20).*

| ***Characteristic*** | ***n*** | ***% of sample*** |
| --- | --- | --- |
| *Gender*  *Female*  *Male* | *6*  *12* | *33.3*  *66.7* |
| *Race*  *Asian*  *Black or African American*  *White*  *Other* | *7*  *1*  *9*  *1* | *38.9*  *5.6*  *50.0*  *5.6* |
| *Handedness*  *Left*  *Right* | *1*  *17* | *5.6*  *94.4* |
| *Highest education level*  *Some college*  *2-year college*  *4-year college*  *Pursuing post graduate degree*  *Post graduate degree* | *2*  *1*  *8*  *2*  *5* | *11.1*  *5.6*  *44.4*  *11.1*  *27.8* |

*Table S2 - Mean intensity for each stimulation condition*

| ***Stimulation Intensity*** | ***Mean (% MSO)*** | ***SD (% MSO)*** | ***Lowest (% MSO)*** | ***Highest (% MSO)*** |
| --- | --- | --- | --- | --- |
| ***Low (110% rMT)*** | *58.42* | *5.90* | *48* | *68* |
| ***High (120-140% rMT)*** | *70.32* | *7.97* | *59* | *84* |

*Table S3 - Intensity of each stimulation condition across participants (N = 18)*

| ***Participant ID*** | ***% MSO of rMT*** | ***Highest Intensity % of rMT*** | ***% MSO for highest intensity*** | ***Lowest Intensity % of rMT*** | ***% MSO for Lowest intensity*** |
| --- | --- | --- | --- | --- | --- |
| *P01* | *51* | *140* | *71* | *110* | *56* |
| *P02* | *51* | *140* | *71* | *110* | *56* |
| *P03* | *56* | *140* | *79* | *110* | *62* |
| *P04* | *50* | *120* | *60* | *110* | *55* |
| *P05* | *55* | *140* | *77* | *110* | *61* |
| *P06* | *56* | *140* | *78* | *110* | *62* |
| *P07* | *NA* | *110* | *NA* | *110* | *NA* |
| *P08* | *61* | *110* | *67* | *110* | *67* |
| *P09* | *48* | *140* | *67* | *110* | *53* |
| *P10* | *47* | *130* | *61* | *110* | *52* |
| *P11* | *45* | *130* | *59* | *110* | *50* |
| *P12* | *48* | *130* | *62* | *110* | *53* |
| *P13* | *44* | *140* | *62* | *110* | *48* |
| *P15* | *60* | *140* | *84* | *110* | *66* |
| *P16* | *55* | *130* | *72* | *110* | *61* |
| *P17* | *50* | *140* | *70* | *110* | *55* |
| *P18* | *55* | *140* | *77* | *110* | *61* |
| *P19* | *62* | *130* | *81* | *110* | *68* |

*Table S4- Timing of each condition in the optimization procedure for all subjects (minutes)*

| ***Participant ID*** | ***Angle optimization in minutes*** | ***Spatial optimization 1.5 cm around the target*** | ***Refining angle*** | ***Sum*** |
| --- | --- | --- | --- | --- |
| *P01* | *12.9* | *10.7* | *10.1* | *33.7* |
| *P02* | *10.2* | *4.1* | *4.1* | *18.4* |
| *P03* | *4.3* | *5.3* | *4.2* | *13.8* |
| *P04* | *7.6* | *6.0* | *4.9* | *18.4* |
| *P05* | *13.5* | *5.9* | *2.1* | *21.5* |
| *P06* | *6.5* | *4.3* | *4.6* | *15.4* |
| *P09* | *1.2* | *4.6* | *2.4* | *8.2* |
| *P10* | *10.9* | *11.6* | *N/A* | *22.5* |
| *P11* | *10.6* | *5.7* | *4.1* | *20.4* |
| *P12* | *12.2* | *3.8* | *4.6* | *20.6* |
| *P13* | *13.0* | *6.4* | *4.9* | *24.4* |
| *P15* | *6.5* | *4.4* | *4.0* | *15.0* |
| *P16* | *7.4* | *4.9* | *2.3* | *14.6* |
| *P17* | *12.1* | *9.0* | *9.1* | *30.3* |
| *P18* | *14.8* | *11.0* | *11.4* | *37.2* |
| *P19* | *13.1* | *8.8* | *12.2* | *34.2* |
| ***Average*** | ***9.8*** | ***6.7*** | ***5.7*** | ***21.8*** |
| ***SD*** | ***3.8*** | ***2.7*** | ***3.3*** | ***8.3*** |

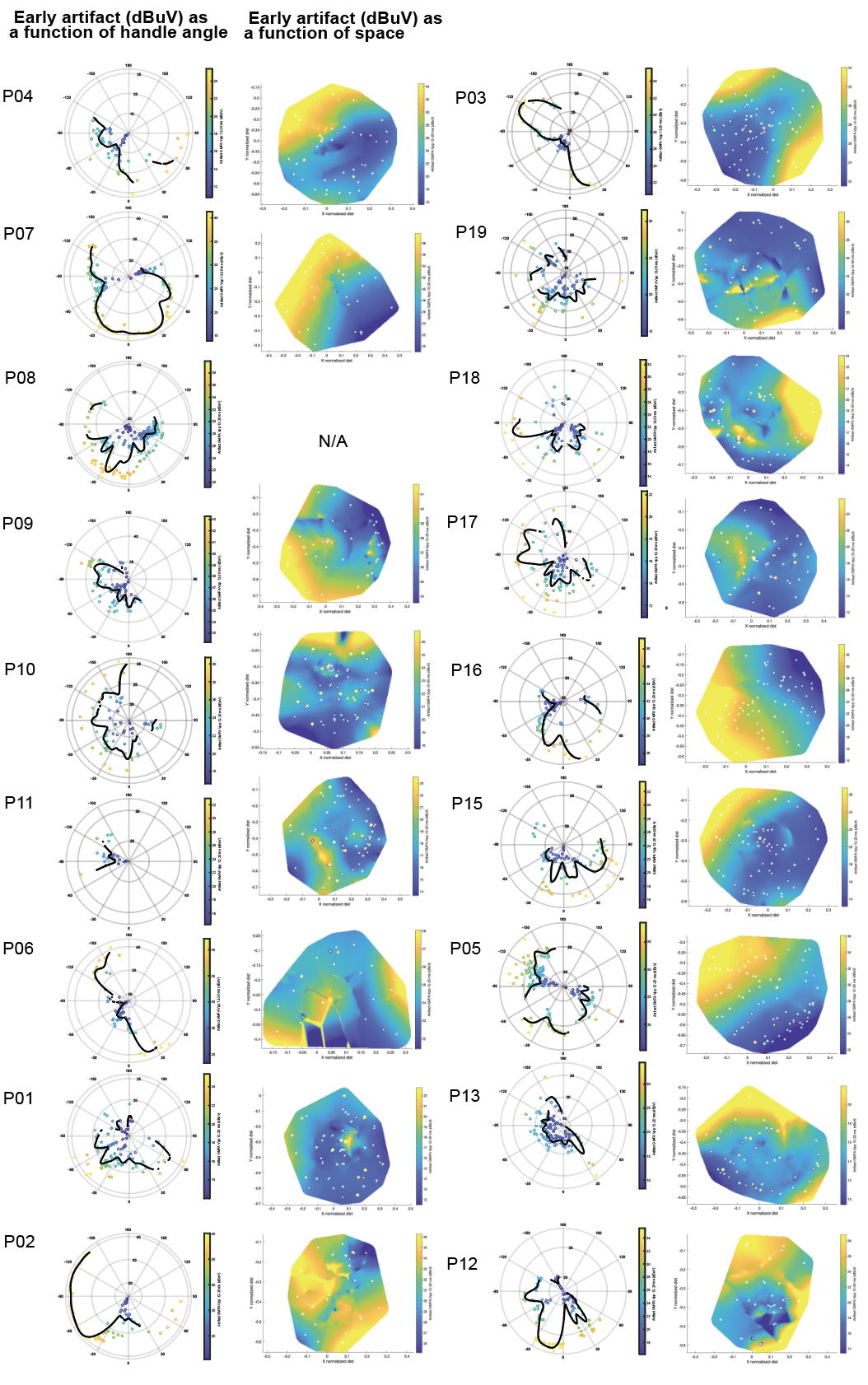

***Fig. S1 - Online angle and location procedure for each participant (n=18) in the dlPFC.*** *Left of each single participant's panel: coil angle search procedure. GMFA of the early muscle artifact (in dBμV at 12-20 ms after TMS) are shown in relation with angles from the midline. Right side of each single participant panel: spatial search for location optimization. GMFA of early artifacts are shown in relation with normalized lateral-medial distance (x-axes) and normalized posterior-anterior distance (y-axes).*

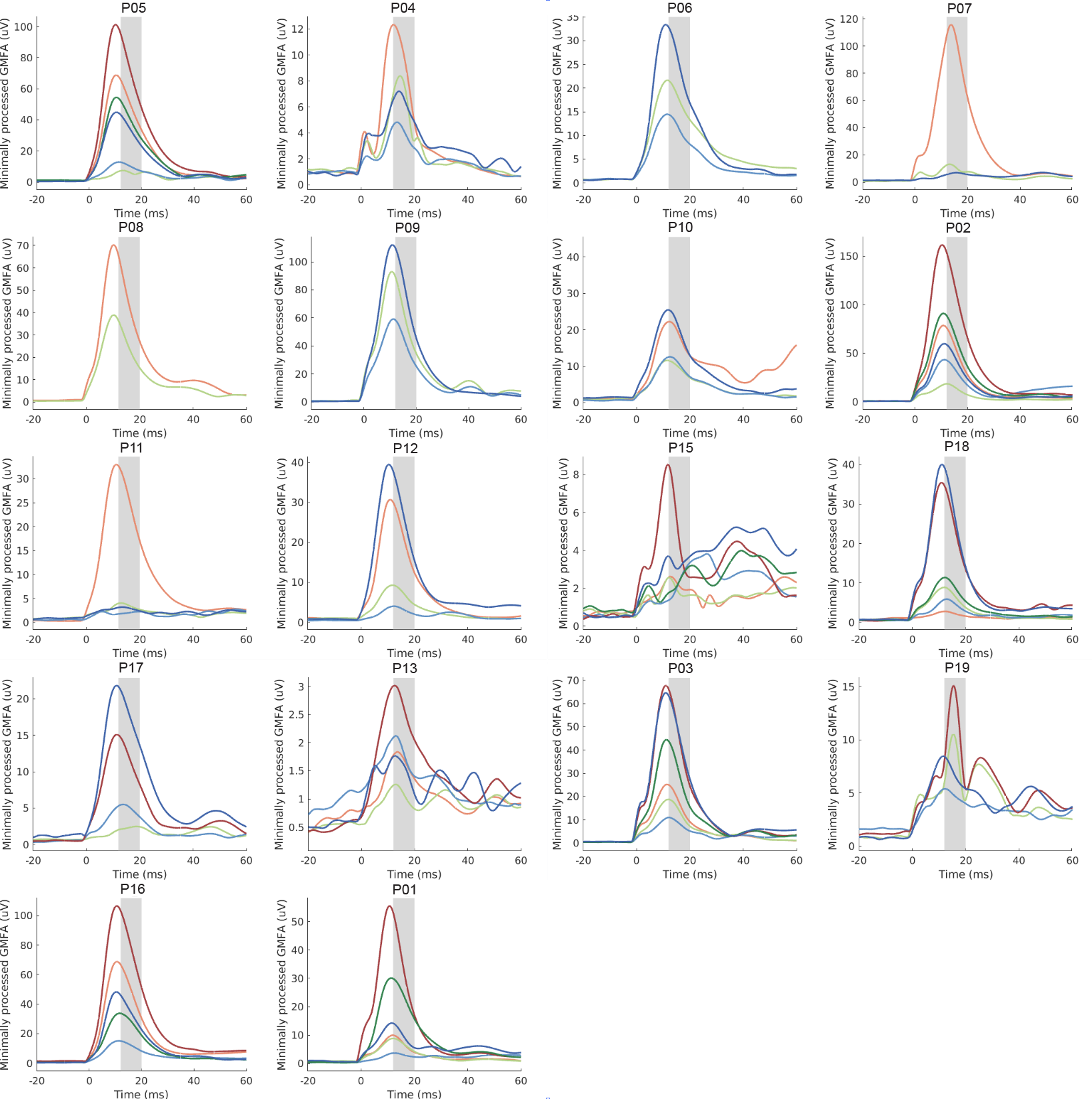

***Fig. S2 - Individual participant early artifact GMFA across conditions.***
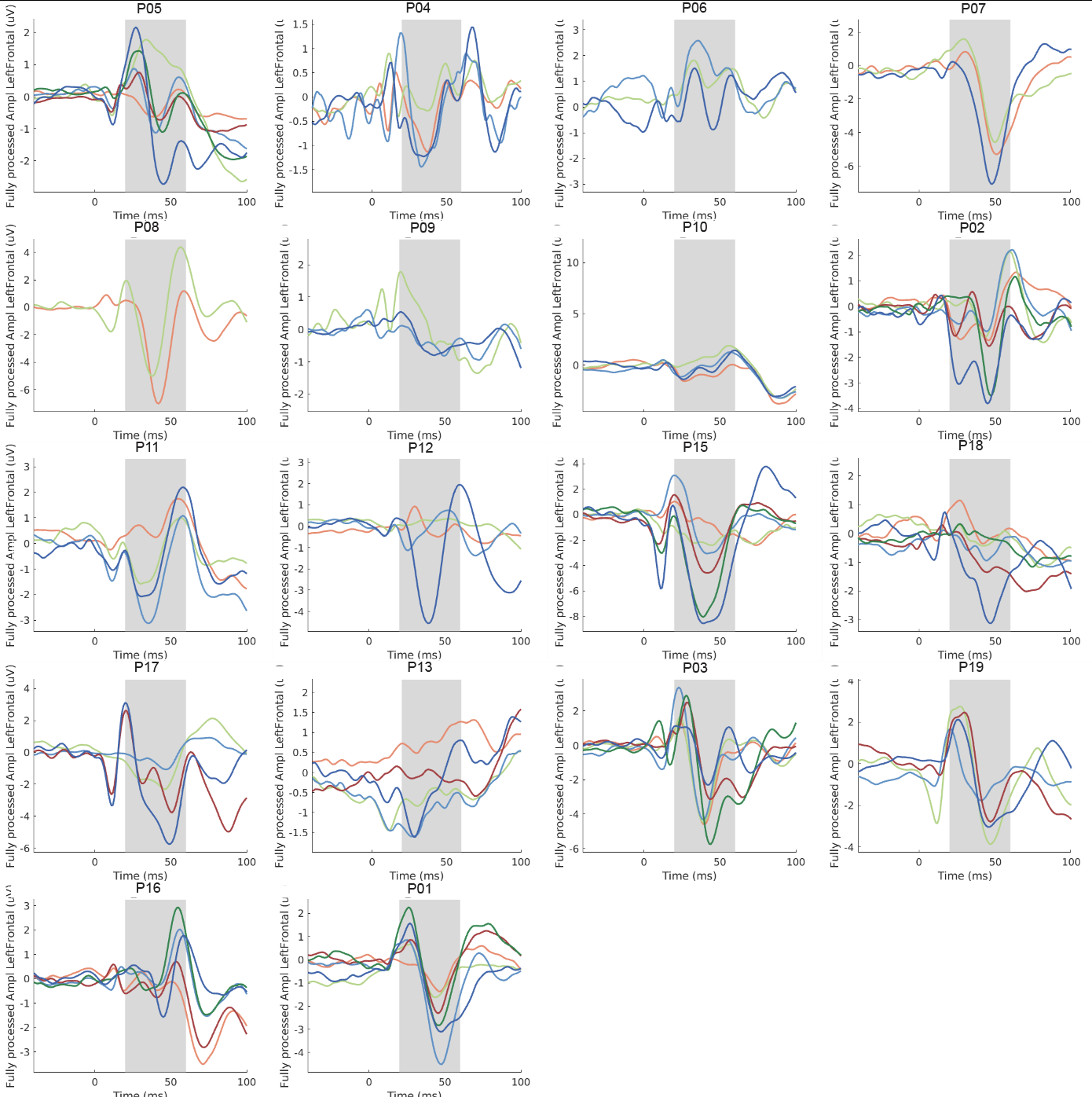

***Fig. S3 - Individual participant local TEP across conditions.*** Data extracted from left frontal ROI.

***Fig. S4 - All participants distal/network responses, A:*** right frontal ROI and ***B:*** left parietal ROI, fully preprocessed data.

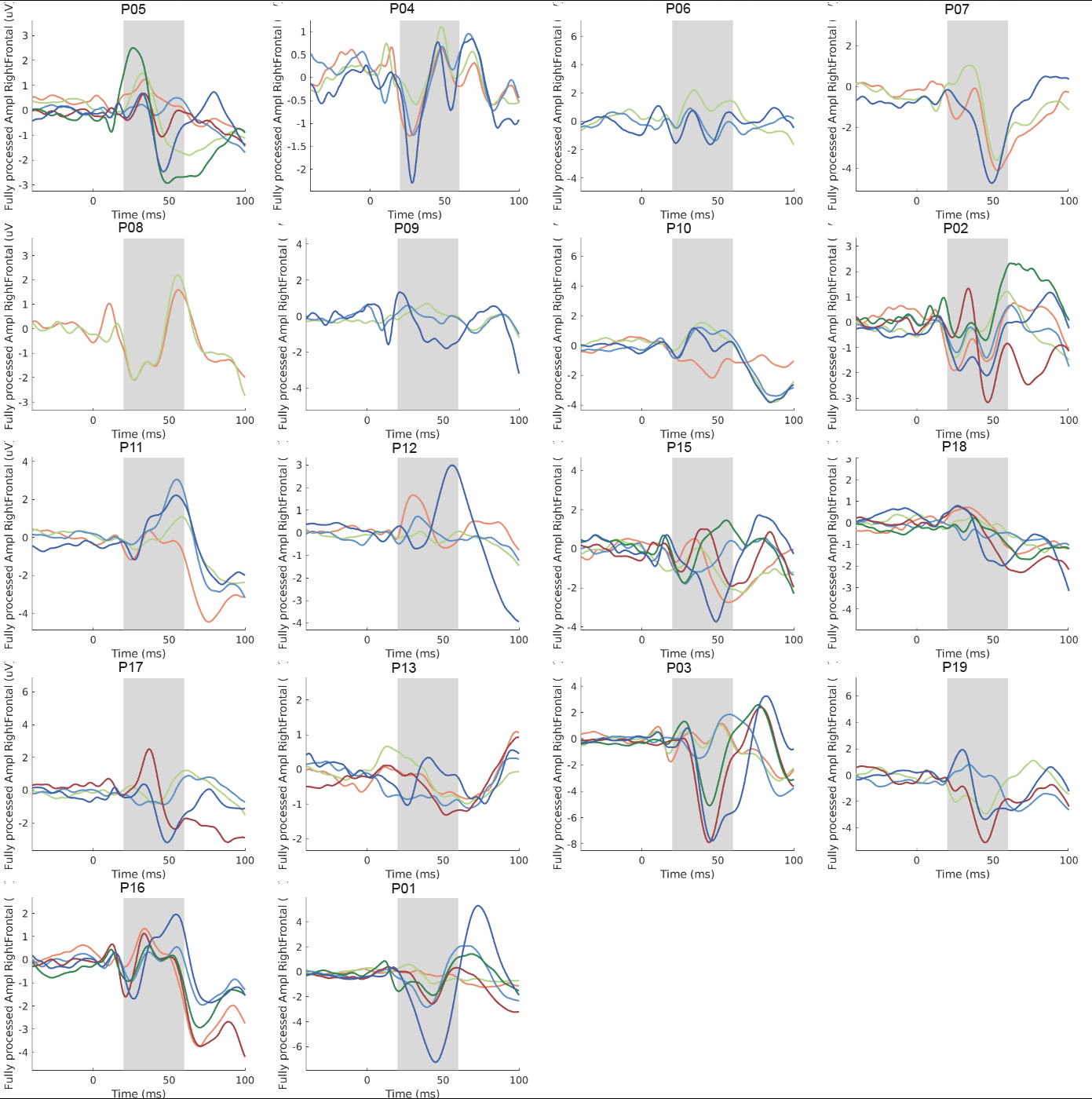

***B:*** left parietal ROI.

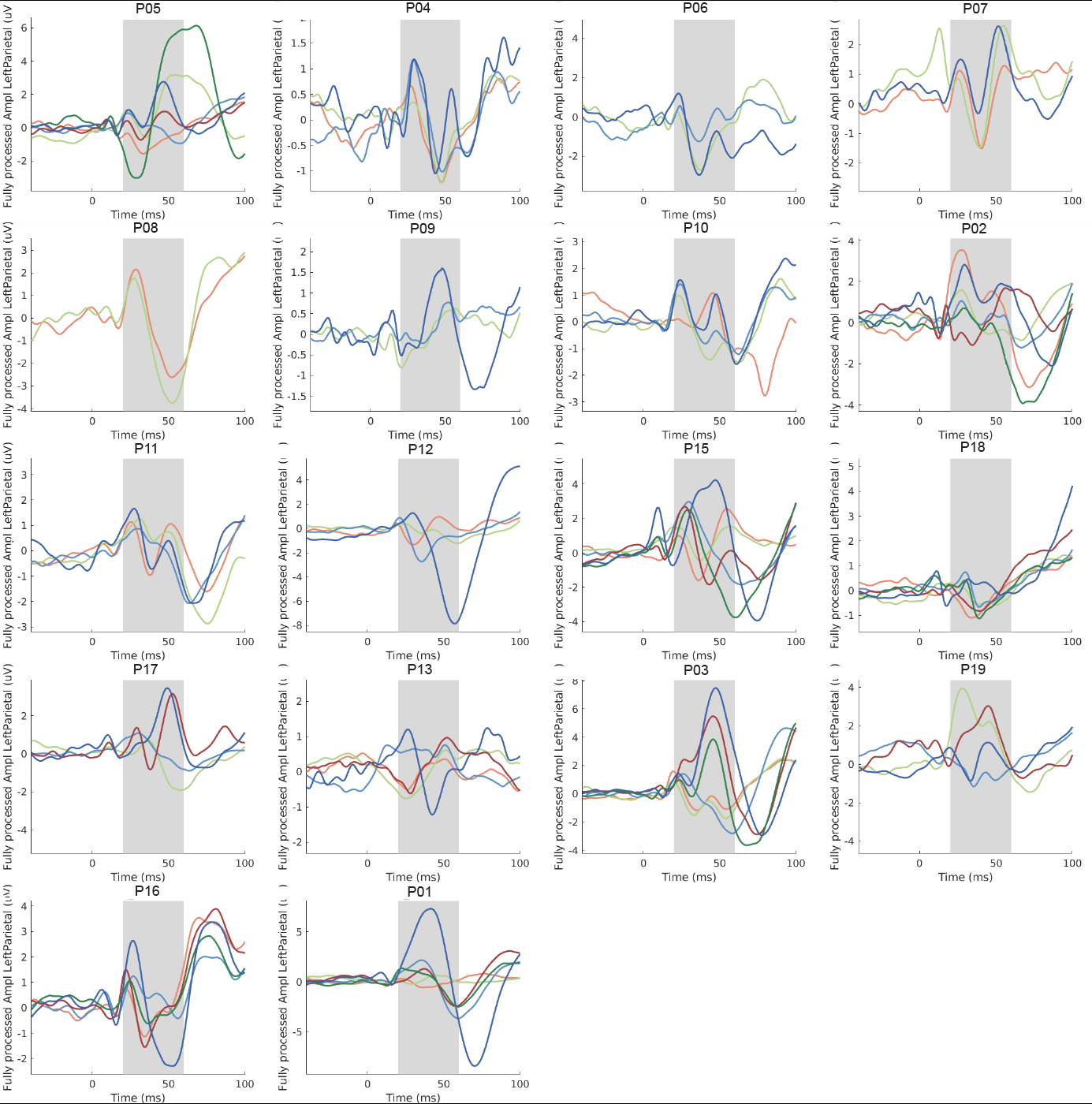

*Table S5- LME results and all EMMeans - Tukey Corrected p-values:* Degrees-of-freedom method: Kenward-Roger. Confidence level used: 0.95

1) **Early Artifact LME**

| ***Optimization*** | ***Intensity*** | ***emmean*** | ***SE*** | ***df*** | ***lower.CL*** | ***upper.CL*** |
| --- | --- | --- | --- | --- | --- | --- |
| Non-optimized | 110% RMT | 23.4 | 2.26 | 26.1 | 18.7 | 28.0 |
| Angle optimized | 110% RMT | 18.0 | 2.28 | 27.0 | 13.3 | 22.6 |
| Angle and location optimized | 110% RMT | 16.7 | 2.24 | 25.6 | 12.1 | 21.3 |
| Non-optimized | highest intensity | 29.5 | 2.53 | 38.1 | 24.3 | 34.6 |
| Angle optimized | highest intensity | 24.9 | 2.75 | 48.6 | 19.4 | 30.5 |
| Angle and location optimized | highest intensity | 23.7 | 2.28 | 27.0 | 19.1 | 28.4 |

2) **Early Artifact Contrasts**

| **contrast** | **estimate** | **SE** | **df** | **t.ratio** | **p.value** |
| --- | --- | --- | --- | --- | --- |
| Non-optimized 110% RMT - Angle optimized 110% RMT | 5.4227458 | 1.637192 | 65.05374 | 3.3122237 | 0.0181267 |
| Non-optimized 110% RMT - Angle and location optimized 110% RMT | 6.6683063 | 1.592417 | 65.06622 | 4.1875368 | 0.0011739 |
| Non-optimized 110% RMT - Non-optimized highest intensity | -6.0821551 | 1.967997 | 65.69774 | -3.0905302 | 0.0333392 |
| Non-optimized 110% RMT - Angle optimized highest intensity | -1.5677814 | 2.251565 | 65.88910 | -0.6963075 | 0.9817444 |
| Non-optimized 110% RMT - Angle and location optimized highest intensity | -0.3627249 | 1.638219 | 65.07773 | -0.2214142 | 0.9999234 |
| Angle optimized 110% RMT - Angle and location optimized 110% RMT | 1.2455605 | 1.620095 | 65.10553 | 0.7688192 | 0.9717862 |
| Angle optimized 110% RMT - Non-optimized highest intensity | -11.5049009 | 1.999692 | 65.81489 | -5.7533373 | 0.0000036 |
| Angle optimized 110% RMT - Angle optimized highest intensity | -6.9905272 | 2.285210 | 66.02182 | -3.0590306 | 0.0362120 |
| Angle optimized 110% RMT - Angle and location optimized highest intensity | -5.7854707 | 1.666613 | 65.13631 | -3.4713946 | 0.0114156 |
| Angle and location optimized 110% RMT - Non-optimized highest intensity | -12.7504614 | 1.962925 | 65.83936 | -6.4956443 | 0.0000002 |
| Angle and location optimized 110% RMT - Angle optimized highest intensity | -8.2360877 | 2.247209 | 65.99917 | -3.6650305 | 0.0063131 |
| Angle and location optimized 110% RMT - Angle and location optimized highest intensity | -7.0310312 | 1.631997 | 65.27488 | -4.3082367 | 0.0007760 |
| Non-optimized highest intensity - Angle optimized highest intensity | 4.5143737 | 2.419078 | 65.29299 | 1.8661546 | 0.4319017 |
| Non-optimized highest intensity - Angle and location optimized highest intensity | 5.7194302 | 1.982488 | 65.63367 | 2.8849754 | 0.0568193 |
| Angle optimized highest intensity - Angle and location optimized highest intensity | 1.2050565 | 2.264157 | 65.83640 | 0.5322318 | 0.9946496 |

3) **Left Frontal EL-TEP LME**

| ***Optimization*** | ***Intensity*** | ***emmean*** | ***SE*** | ***df*** | ***lower.CL*** | ***upper.CL*** |
| --- | --- | --- | --- | --- | --- | --- |
| Non-optimized | 110% RMT | 6.36 | 1.48 | 37.5 | 3.37 | 9.35 |
| Angle optimized | 110% RMT | 5.52 | 1.50 | 39.3 | 2.48 | 8.55 |
| Angle and location optimized | 110% RMT | 8.57 | 1.46 | 36.3 | 5.60 | 11.53 |
| Non-optimized | highest intensity | 7.50 | 1.77 | 58.6 | 3.96 | 11.04 |
| Angle optimized | highest intensity | 10.85 | 2.01 | 71.0 | 6.85 | 14.85 |
| Angle and location optimized | highest intensity | 11.83 | 1.50 | 39.3 | 8.79 | 14.86 |

4) **Left Frontal EL-TEP Contrasts**

| **contrast** | **estimate** | **SE** | **df** | **t.ratio** | **p.value** |
| --- | --- | --- | --- | --- | --- |
| Non-optimized 110% RMT - Angle optimized 110% RMT | 0.8434254 | 1.420074 | 65.12248 | 0.5939305 | 0.9911029 |
| Non-optimized 110% RMT - Angle and location optimized 110% RMT | -2.2052343 | 1.381178 | 65.14510 | -1.5966332 | 0.6037010 |
| Non-optimized 110% RMT - Non-optimized highest intensity | -1.1392265 | 1.703467 | 66.47595 | -0.6687695 | 0.9847605 |
| Non-optimized 110% RMT - Angle optimized highest intensity | -4.4869118 | 1.947755 | 66.90366 | -2.3036331 | 0.2071407 |
| Non-optimized 110% RMT - Angle and location optimized highest intensity | -5.4669181 | 1.420851 | 65.16924 | -3.8476370 | 0.0035828 |
| Angle optimized 110% RMT - Angle and location optimized 110% RMT | -3.0486597 | 1.405012 | 65.23168 | -2.1698463 | 0.2658895 |
| Angle optimized 110% RMT - Non-optimized highest intensity | -1.9826519 | 1.730243 | 66.72179 | -1.1458808 | 0.8602434 |
| Angle optimized 110% RMT - Angle optimized highest intensity | -5.3303373 | 1.976009 | 67.18398 | -2.6975262 | 0.0891143 |
| Angle optimized 110% RMT - Angle and location optimized highest intensity | -6.3103436 | 1.445209 | 65.29514 | -4.3663885 | 0.0006348 |
| Angle and location optimized 110% RMT - Non-optimized highest intensity | 1.0660078 | 1.698297 | 66.77407 | 0.6276923 | 0.9885573 |
| Angle and location optimized 110% RMT - Angle optimized highest intensity | -2.2816775 | 1.943291 | 67.13537 | -1.1741304 | 0.8475100 |
| Angle and location optimized 110% RMT - Angle and location optimized highest intensity | -3.2616838 | 1.414557 | 65.58327 | -2.3057987 | 0.2065622 |
| Non-optimized highest intensity - Angle optimized highest intensity | -3.3476853 | 2.096695 | 65.64592 | -1.5966486 | 0.6036693 |
| Non-optimized highest intensity - Angle and location optimized highest intensity | -4.3276916 | 1.716366 | 66.34145 | -2.5214266 | 0.1326877 |
| Angle optimized highest intensity - Angle and location optimized highest intensity | -0.9800063 | 1.958983 | 66.79293 | -0.5002628 | 0.9960010 |

5) **Right Frontal TEP LME**

| **Optimization** | **Intensity** | **emmean** | **SE** | **df** | **lower.CL** | **upper.CL** |
| --- | --- | --- | --- | --- | --- | --- |
| Non-optimized | 110% RMT | 5.29 | 1.22 | 44.4 | 3.18 | 8.08 |
| Angle optimized | 110% RMT | 5.29 | 1.24 | 46.5 | 2.80 | 7.78 |
| Angle and location optimized | 110% RMT | 5.38 | 1.20 | 42.8 | 2.96 | 7.80 |
| Non-optimized | highest intensity | 10.60 | 1.50 | 66.9 | 7.61 | 13.59 |
| Angle optimized | highest intensity | 9.28 | 1.72 | 76.9 | 5.86 | 12.70 |
| Angle and location optimized | highest intensity | 10.98 | 1.24 | 46.5 | 8.48 | 13.47 |

6) **Right Frontal TEP Contrasts**

| **contrast** | **estimate** | **SE** | **df** | **t.ratio** | **p.value** |
| --- | --- | --- | --- | --- | --- |
| Non-optimized 110% RMT - Angle optimized 110% RMT | 0.3375908 | 1.275933 | 65.16822 | 0.2645835 | 0.9998158 |
| Non-optimized 110% RMT - Angle and location optimized 110% RMT | 0.2492490 | 1.240952 | 65.19483 | 0.2008530 | 0.9999527 |
| Non-optimized 110% RMT - Non-optimized highest intensity | -4.9679452 | 1.528600 | 66.93791 | -3.2499963 | 0.0213975 |
| Non-optimized 110% RMT - Angle optimized highest intensity | -3.6484161 | 1.747158 | 67.51797 | -2.0882007 | 0.3058067 |
| Non-optimized 110% RMT - Angle and location optimized highest intensity | -5.3453944 | 1.276568 | 65.22636 | -4.1873149 | 0.0011722 |
| Angle optimized 110% RMT - Angle and location optimized 110% RMT | -0.0883419 | 1.262271 | 65.31133 | -0.0699865 | 0.9999998 |
| Angle optimized 110% RMT - Non-optimized highest intensity | -5.3055360 | 1.552263 | 67.25888 | -3.4179374 | 0.0131814 |
| Angle optimized 110% RMT - Angle optimized highest intensity | -3.9860070 | 1.772029 | 67.88513 | -2.2494025 | 0.2294430 |
| Angle optimized 110% RMT - Angle and location optimized highest intensity | -5.6829852 | 1.298304 | 65.39333 | -4.3772370 | 0.0006105 |
| Angle and location optimized 110% RMT - Non-optimized highest intensity | -5.2171941 | 1.523528 | 67.32793 | -3.4244160 | 0.0129273 |
| Angle and location optimized 110% RMT - Angle optimized highest intensity | -3.8976651 | 1.742768 | 67.82095 | -2.2364794 | 0.2350566 |
| Angle and location optimized 110% RMT - Angle and location optimized highest intensity | -5.5946433 | 1.270417 | 65.76758 | -4.4037833 | 0.0005534 |
| Non-optimized highest intensity - Angle optimized highest intensity | 1.3195291 | 1.882998 | 65.86980 | 0.7007596 | 0.9812199 |
| Non-optimized highest intensity - Angle and location optimized highest intensity | -0.3774492 | 1.540373 | 66.76222 | -0.2450375 | 0.9998739 |
| Angle optimized highest intensity - Angle and location optimized highest intensity | -1.6969783 | 1.757416 | 67.37330 | -0.9656098 | 0.9271438 |

7) **Left Parietal LME**

| **Optimization** | **Intensity** | **emmean** | **SE** | **df** | **lower.CL** | **upper.CL** |
| --- | --- | --- | --- | --- | --- | --- |
| Non-optimized | 110% RMT | 7.07 | 1.16 | 49.2 | 4.74 | 9.39 |
| Angle optimized | 110% RMT | 6.80 | 1.18 | 51.5 | 4.43 | 9.17 |
| Angle and location optimized | 110% RMT | 7.02 | 1.14 | 47.3 | 4.72 | 9.31 |
| Non-optimized | highest intensity | 9.60 | 1.45 | 71.2 | 6.72 | 12.49 |
| Angle optimized | highest intensity | 11.62 | 1.68 | 79.2 | 8.29 | 14.96 |
| Angle and location optimized | highest intensity | 10.71 | 1.18 | 51.5 | 8.33 | 13.08 |

8) **Left Parietal Contrasts**

| **contrast** | **estimate** | **SE** | **df** | **t.ratio** | **p.value** |
| --- | --- | --- | --- | --- | --- |
| Non-optimized 110% RMT - Angle optimized 110% RMT | 0.2670370 | 1.274374 | 65.20380 | 0.2095437 | 0.9999417 |
| Non-optimized 110% RMT - Angle and location optimized 110% RMT | 0.0499254 | 1.239412 | 65.23228 | 0.0402816 | 1.0000000 |
| Non-optimized 110% RMT - Non-optimized highest intensity | -2.5364170 | 1.525281 | 67.27171 | -1.6629177 | 0.5606428 |
| Non-optimized 110% RMT - Angle optimized highest intensity | -4.5576029 | 1.742876 | 67.96766 | -2.6149904 | 0.1075735 |
| Non-optimized 110% RMT - Angle and location optimized highest intensity | -3.6396753 | 1.274962 | 65.26911 | -2.8547319 | 0.0613327 |
| Angle optimized 110% RMT - Angle and location optimized 110% RMT | -0.2171115 | 1.260632 | 65.37134 | -0.1722244 | 0.9999780 |
| Angle optimized 110% RMT - Non-optimized highest intensity | -2.8034540 | 1.548621 | 67.64626 | -1.8102904 | 0.4662089 |
| Angle optimized 110% RMT - Angle optimized highest intensity | -4.8246399 | 1.767334 | 68.39697 | -2.7298970 | 0.0823436 |
| Angle optimized 110% RMT - Angle and location optimized highest intensity | -3.9067123 | 1.296560 | 65.46631 | -3.0131362 | 0.0409537 |
| Angle and location optimized 110% RMT - Non-optimized highest intensity | -2.5863425 | 1.519899 | 67.72755 | -1.7016540 | 0.5355278 |
| Angle and location optimized 110% RMT - Angle optimized highest intensity | -4.6075284 | 1.738210 | 68.32153 | -2.6507316 | 0.0990885 |
| Angle and location optimized 110% RMT - Angle and location optimized highest intensity | -3.6896007 | 1.268452 | 65.90149 | -2.9087431 | 0.0534761 |
| Non-optimized highest intensity - Angle optimized highest intensity | -2.0211859 | 1.880044 | 66.03881 | -1.0750737 | 0.8895221 |
| Non-optimized highest intensity - Angle and location optimized highest intensity | -1.1032582 | 1.537175 | 67.06659 | -0.7177180 | 0.9791363 |
| Angle optimized highest intensity - Angle and location optimized highest intensity | 0.9179276 | 1.753247 | 67.79877 | 0.5235587 | 0.9950515 |
